# How complexity originates: The evolution of animal eyes

**DOI:** 10.1101/017129

**Authors:** Todd H. Oakley, Daniel I. Speiser

## Abstract

Learning how complex traits like eyes originate is fundamental for understanding evolution. Here, we first sketch historical perspectives on trait origins and argue that new technologies offer key new insights. Next, we articulate four open questions about trait origins. To address them, we define a research program to break complex traits into components and study the individual evolutionary histories of those parts. By doing so, we can learn when the parts came together and perhaps understand why they stayed together. We apply the approach to five structural innovations critical for complex eyes, reviewing the history of the parts of each of those innovations. Photoreceptors evolved within animals by bricolage, recombining genes that originated far earlier. Multiple genes used in eyes today had ancestral roles in stress responses. We hypothesize that photo-stress could have increased the chance those genes were expressed together in places on animals where light was abundant.

## 1. Introduction

How do complex (multi-part and functionally integrated) biological traits such as eyes, feathers and flight, metabolic pathways, or flowers, originate during evolution? These biological features often appear so functionally integrated and so complicated, that imagining the evolutionary paths to such complexity is sometimes difficult. While it is clear that structurally and functionally complex systems originated through evolutionary processes, broad questions still remain about which evolutionary processes more commonly lead to innovation and complexity. Here, we use eye evolution as a focus for how to implement a research program to gain understanding of the origin of complex traits. This research program first requires defining the trait in question, followed by inferring with comparative methods the timing of past evolutionary events including changes in function. By understanding *when* different components came together, we can begin to understand *how* they came together, and by making inferences about possible functions and environmental context, we can begin to understand *why* those components stayed together.Eye evolution is particularly amenable to such a research program because we can use optics to predict function from morphology and we can use extensive knowledge about the genetic components of eyes and light sensitivity to predict and test gene functions in a broad range of organisms. This approach leads to a narrative on animal eye evolution that, while still incomplete, is already rich and detailed in many facets. We know that a diversity of sometimes disparate eyes evolved using functional components that interact with light. All of these components are used outside of eyes and all were recruited into light receptive organs during evolution at many different times and in many different combinations. One common theme that emerges is that many eye-genes had ancestral roles related to stress response. Therefore, evolved responses to light as a stressor may have brought together many of the genes that today function in eyes.

## 2. Past and present perspectives on origins

Because they still influence many people’s understanding of evolution, we begin with a brief sketch of two historical perspectives on trait origins. One we call a gradual-morphological perspective, the other we call a binary phylogenetic perspective. We argue that these two historical perspectives were incomplete, or even misleading. The gradual-morphological perspective simply assumes variation exists, without considering its source. The binary phylogenetic perspective makes the oversimplifying assumption that all components of traits are gained and lost as one. In this section, we explain these perspectives, their shortcomings, and then explain how new information and new technology allows us to enrich our understanding of trait origins compared to these historical perspectives.

### a. Two incomplete perspectives on trait origins

One perspective, which we call the ‘gradual-morphological model’, provides a typical evolutionary narrative for the origins of complex traits. The model involves gradual elaboration driven by natural selection (e.g. 1–3). When applied to eyes, the model begins with a light sensitive patch of cells. That patch evolves to form a deeper and deeper cup, and finally an increasingly more efficient lens evolves within the cup. Such a gradual progression from simple light sensitive patch to complex eye was first imagined by Darwin (2) when he explained a corollary to his new biological mechanism of natural selection: If complex traits like eyes were produced by natural selection, there should exist a series of functional intermediates between simple patch and complex eye. To show this condition is met, Darwin mentioned functional variation in eye complexity in different species. Later, Salvini-Plawen and Mayr (1) illustrated several cases where related species, such as various snails, showed rather finely graded variation between eye-spot and lens-eye. Taking the idea one step farther, Nilsson and Pelger (3) quantified the gradual morphological changes probably necessary to evolve from patch to eye, and estimated that the progression can occur rapidly in geological time. One commonly sees similar variations on the gradual-morphological model in textbooks and in popular books and videos to help explain how natural selection could have produced an eye.

While this gradual-morphological progression is logical and provides a powerful and visual way to imagine the stepwise evolution of complexity, it also has at least two shortcomings (4). First, the linear series of eyes arranged from simple to complex implies incorrectly that evolution always proceeds that way (4). Yet evolution often results in loss or reduced functionality of structures, including eyes (5). Second, in the gradual models, variation is simply assumed without being addressed. In fact, each gradual step along the progression requires natural selection to act upon variation. However, the developmental-genetic basis for how this morphological variation originates was not considered (understandably, given the technological limitations of the time). Further, discrete origins were not considered, except again through assuming that the variation simply arose in the past. For example, the origin of light sensitivity itself was usually not addressed, nor was the origin of the cup structure, or the origin of the first lens material. These discrete origins of photosensitivity, of a depression, or of lens material were treated no differently than the gradual elaboration of existing structures. Therefore, while not uninformative, using a gradual series of eyes as a model for how evolution proceeds was incomplete, in that it assumed morphological variation without addressing the mechanisms leading to variation. How did light sensitivity originate? How did lenses or eye-pigmentation originate? Understanding these are critical for a complete picture of eye origins and evolution. Herein we will suggest comparative approaches to these questions.

Another approach, which we call a binary-phylogenetics, makes some potentially misleading assumptions. By focusing on the distribution of complex traits in different species, phylogeneticists often gain important insights into the timing of evolutionary events. But when they score such traits as simply absent or present, they cannot gain insights into *how* traits originated because they implicitly assume all components of the trait evolve in concert. For example, armed with a phylogenetic tree, an evolutionist might score species as ‘eyed’ or ‘eyeless’, to infer the number and/or timing of eye gains and losses (e.g. 6; Fig. 1). Based on assumptions like parsimony or maximum likelihood, one may then suggest that character states scored in living species can be inferred in common ancestors, leading to inferences of trait history as series of all-or-none gains and losses. Even if the separate components of multi-part systems have different evolutionary histories, scoring complex traits as simply present or absent makes inferring those separate histories impossible. The inference of all-or-none gains and losses can be said to impose a punctuated mode of evolution, such that all components of a complex trait originate or become extinct simultaneously (but see reference 7 for an alternative phylogenetic model).

**Figure 1. A.**
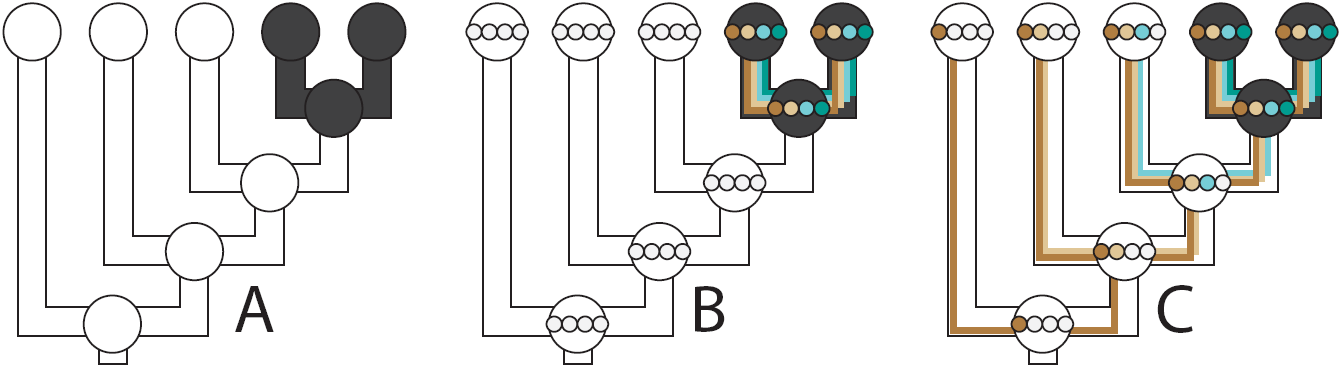
Phylogeneticists sometimes score complex traits like eyes as absent (white) or present (black) in different species. This makes the implicit assumption that all components of that trait share the same evolutionary history. **B**. However, complex traits comprise many components, as illustrated by four small, colored circles. For eyes, these might be lenses, opsin genes, pigments, and ion channels. By only scoring complex traits as present or absent, we force a punctuated mode of evolution, whereby all components are gained and lost together. Under such a model, a complex trait and each of its components is either fully present or fully absent. **C**. Illustrates a more gradual mode of component evolution, where one component is added at each of 4 different ancestral nodes. Explicitly considering the histories of the components, could allow inference of this gradual mode of evolution.

### b. New information, new perspectives on origins

New information and new technologies allow for new perspectives on evolutionary origins of traits. Perhaps the biggest advances in understanding evolutionary origins come from our ever increasing understanding of connections between genotype and phenotype, fueled by knowledge about the molecular components of traits. Historically, it was not possible to go far beyond the gradual-morphological model of eye evolution because understanding the developmental-genetic basis of variation in traits was not possible. Information was also missing to extend comparative perspectives on origins. While phylogeneticists historically could score the presence or absence of a trait like an eye by simply looking at a species, knowing the components of those eyes and their separate histories requires more information. Even when molecular components became known from a model organism, extending that knowledge outside models was not feasible, making comparative, evolutionary studies intractable. We now know many molecular components of eyes, which we review below, and we now feasibly can obtain extensive information about these components from non-model organisms. When these components are proteins, as is often the case, we can trace their individual evolutionary histories. Instead of forcing the all-or-none perspective that is implicit in scoring traits like eyes as ‘present’ or ‘absent’, tracing individual histories of components allows us to understand that some components are ancient and others are new. In this way, understanding when components of multi-part systems came together is leading us to new questions and insights about evolutionary origins.

## 3. Four open questions on origins: eye evolution and beyond

### a. Do multi-part traits form gradually or abruptly?

The different parts of complex traits like eyes could come together gradually, where they are added sequentially over longer periods of time, or could originate abruptly, with all the parts of a trait coming together in a short period of time (8). Gradual versus punctuated patterns of origin are two ends of a spectrum, and intermediates are also possible: A particular trait may have had bouts of both gradual and punctuated addition of components before arriving at its current state. Furthermore, some traits may have originated gradually and others abruptly. Armed with increasing knowledge of traits’ components, and with phylogenetic methods to reconstruct the timing of their origins, we are now in a position to elucidate the origins of eyes and other traits to determine gradual versus punctuated origins, which could lead to a general understanding of the mechanisms that lead to the origin of complex traits. Similar ideas were explored by Plachetzki and Oakley (8), where they suggested that co-duplication of all parts of a trait (punctuated change) could serve as a null model for the origin of multi-part systems. The inferred origin(s) of traits can serve as a null expectation for the timing of origin of each part. If parts are older than the trait, then those parts may have been co-opted or recruited from a previously existing function into the newly evolved trait. Co-duplication serves as a useful null expectation because it can be rejected by any part of a complex trait. The dual phototransduction systems of vertebrate retinas, one used in rods and the other in cones (9, 10), may be a prime example of co-duplication. In contrast, there are few other examples of co-duplication, which is often rejected in favor of co-option (11). One of the reasons that co-option may be much more common than co-duplication is that mutational events required for co-duplication may be less common.

### b. What types of mutations are involved in origins?

Mutations that became fixed in populations are the primary source of evolutionary change, and various types of mutation may be more commonly associated with trait origins than others. Here, we differentiate the mutational requirements for two modes of trait origin - co-duplication and co-option. Co-duplication means that multiple parts of a trait originate simultaneously. Simultaneous duplication requires large scale copying of entire genomes or chromosomal blocks, which did occur early in vertebrate history to lead to the duplication of phototransduction genes (9). Following such block duplication, each set of duplicates must also diverge in function, which requires other mutations after the duplication event. For co-duplication to occur without whole genome duplication (and without a large number of simultaneous, yet independent mutations), co-functioning genes could also be located near each other on a chromosome so they could be copied all at once. While not as obvious in animals as bacteria with operons, a recent study found co-functioning genes to be located non-randomly in the human genome (12). Therefore, the spatial association of co-functioning genes could facilitate origins of new traits by co-duplication.

Co-option is defined as ‘the acquisition of new roles by ancestral characters’ (13), yet often conflates pattern and process or structure and function. Given the diversity of meanings of co-option, a comprehensive survey of the mutational causes is challenging. For the purposes of this review, we simply point out that co-option will often involve regulatory mutations that cause a gene (whether duplicated or not) to be expressed in a new place. Because there is no general relationship between regulatory sequence and spatiotemporal gene expression, inferring the history of co-option can be challenging. Usually, inferring co-option requires comparing gene function to the phylogenetic history of those genes (8, 14). Therefore, the specific mutational source of co-option may often remain unknown, especially in very ancient comparisons.

### c. What originates first, structure or function?

How do functional changes relate to structural changes during evolution (see also 15)? It turns out that this is an enduring yet underexplored question about trait origins (2, 16, 17). One possibility is that structural changes arise first (Fig. 2a). For example, a gene could duplicate first (a structural change in DNA), followed later by a gain in function in one of the new genes. This corresponds to the classic neo-functionalization model of gene evolution (18). Similarly, a cell type, organ, or other structure could duplicate (as defined by 11) during evolution, followed by a change in function in one of the new structures. This neo-functionalization process would yield a pattern whereby closely related structures have different functions, and outgroup structures share a function with one of the ingroup modules (Fig. 2a). Alternatively, functional changes could evolve first. Here, a biological structure like a gene, cell, or organ could first gain a function, making it multi-functional, and sometimes called gene sharing (19). Later, the structure could duplicate and subdivide the ancestral functions between the duplicates, in a subfunctionalization or division of labor mode of evolution (2, 20, 21). Sub-functionalization could yield a pattern different from neo-functionalization, whereby sub-functionalization leads to two closely related modules with different functions and a multifunctional outgroup module (Fig. 2b).

**Figure 2.**
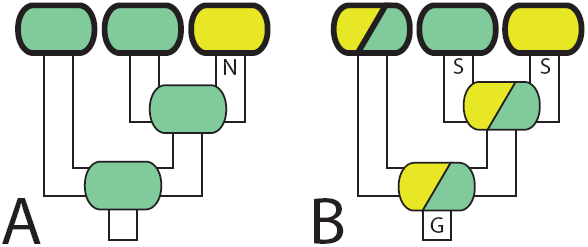
Rounded rectangles represent a generalized biological structure (gene, pathway, organ, etc). Green and blue colors represent different functions, broadly construed (including spatial, temporal, or contextual differences as well as separate functions). The functions at the tips of the trees are observed (illustrated with a bolder outline) but functions at nodes are inferred with phylogenetic techniques. **A**. A new structure evolves first, before a new function. N=Neo-functionalization (equals a gain of function). Here, two closely related “ingroup” structures have distinct functions. The “outgroup” structure has the same function as one of the ingroup members. In this case, parsimony favors a single change in function leading to the functionally unique ingroup structure. In the gene duplication literature, this is termed Neo-Functionalization (20). **B**. A new function evolves first, before the structure duplicates. G=gain of function, S=Subfunctionalization. Here two closely related “ingroup” structures have non-overlapping functions, and an “outgroup” structure performs both functions. With this phylogenetic distribution of structures and functions, a parsimonious explanation is Subfunctionalization or Division of labor, such that an ancestral structure had two functions that specialized after duplication. In both A and B, other evolutionary histories are possible that could lead to concluding the opposite evolutionary process, but these require more events, and are less parsimonious.

### d. What is the environmental context of origins?

Inferring when and where traits and their components originated could give information about the environmental context of origins. Some broad patterns about origins are already evident from paleontological distributions and phylogenetic studies. For example, many evolutionary novelties that define major taxonomic clades originated in the tropics (22) and in shallow marine environments (23). In addition, many trait origins are correlated with organismal changes in environment, such as transitions between aquatic and terrestrial, marine and freshwater, and benthic and pelagic (24) life histories. The origins of image forming eyes may be correlated with transitions to active lifestyles (25). These general patterns relate to species-level environmental interactions, but similar ideas can be applied to other levels, like gene or cell type. For example, genes expressed in different organismal locations experience different cellular environments. One logical hypothesis about cellular environments and origins is that light creates a stressful environment for cells, such that the origins of light interacting genes may often be related to responses to light stress. We return to the light stress origins hypothesis in Sections 5 and 6.

## 4. A Research Program to Investigate Origins

### a. Define the trait

How can we begin to address unresolved questions about origins of multi-part systems like eyes? The first step is to define the trait in question, which involves enumerating its parts (we often use ‘components’ interchangeably with ‘parts’ in this review). Because of the sheer number of parts, highly complex structures like image-forming eyes are difficult to fully enumerate. In particular, eyes contain many generic genetic components, like basic cellular machinery, which will not be especially informative about the evolution of eyes per se. Identifying the parts unique to eyes may sound tempting. However, with this strategy, most (or even all) components would be excluded because they have functions outside eyes, even if those parts are functionally critical to eyes. In addition to methods to analyze entire expression profiles of traits (e.g. 26), another way forward is to identify and explicitly define modules with important functions. For visual systems, approachable modules whose evolutionary histories can be understood include pigment synthesis pathways and phototransduction cascades. We also know of structural components of lenses and corneas, whose histories can be traced. These parts are enumerated and established in model organisms, and we can search for similar components in the rapidly growing database of full genome sequences. Further, high throughput sequencing in non-model organisms can produce detailed transcriptome sequences, where components can be identified by similarity to known genes (27, 28). After defining the trait, we can begin to understand when, how, and why complex traits like eyes originated during evolution.

### b. Estimate when the trait originated

Estimating the relative or absolute timing of trait origins can be done using phylogenetic methods to compare the distribution of presence/absence of traits to phylogenetic history (Fig. 1a). For example, if two species share a trait, we might infer their common ancestor also had that trait, so the trait originated earlier. Specifically for understanding the history of animal eyes and their components, major animal clades become a focus. We will see that many light interaction components are well characterized in protostomes like flies and deuterostomes like vertebrates, indicating an origin of the trait at or before bilaterians. Many traits are present in bilaterians and cnidarians, implying an origin at or before Eumetazoa, and many traits are present also in sponges or even choanoflagellates, implying an origin before animals. This core logic for determining relative timing of a trait compared to a tree is based on parsimony, and the enterprise of understanding the evolutionary history of traits, often called ancestral state reconstruction or character mapping, has developed a rich array of statistical techniques (29).

We can also estimate the absolute timing of trait origins when character mapping is performed on a time-calibrated phylogeny (30). Estimating character histories and the absolute timing of evolutionary events is not trivial, and there is an extensive literature in phylogenetics on the challenges involved, including limitations of character evolution models (e.g. 29) and challenges of accurately estimating divergence times using relaxed molecular clock models and fossil calibrations (e.g. 31). Despite these challenges, estimating when a particular trait first evolved serves as an initial point of comparison for the evolutionary histories of the parts that make up the trait under investigation. While this initial estimate is a starting point, one quickly realizes a major challenge: different components of a trait invariably have different histories. So instead of an all-or-none tracing of traits on a phylogeny, the individual histories of those parts and how they came to function together must be the focus. Therefore, the critical next step is to understand those separate histories.

### c. Determine when the components originated and became functionally associated

Once a trait and target suite of parts is identified - such as the genes used to make a pigment or a lens - a next step is to determine when the parts themselves originated, for genes sometimes termed ‘Phylostratigraphy’ (32). However, similar genetic structures do not always imply similar functions. A full understanding of the history of complex traits also requires estimating the history of function. To understand trait origins, we must consider several aspects of function. First, a gene’s biochemical function is always of critical importance. Second, a gene’s interaction partners are critical to function. These are usually mediated by protein-protein interactions. Third, the site and time of expression of a gene has critical implications for function. Finally, a particular gene is of interest because of how it contributes to an organismal function. All of these functions change over evolutionary time, and because we do not have direct ways to test the function of genes from long extinct ancestors, we must rely on comparative analyses of function in living organisms. Since comparative analyses of function rely on sound experimental demonstrations of function across organisms at different levels of function, understanding the evolutionary origins of complex traits requires a diverse suite of information from genetics, biochemistry, physiology, development, behavior, and phylogenetics. We next summarize the state of this information relating to the evolution of light sensing systems in animals.

## 5. Light Interaction Genes and Their Evolutionary Origins

A recent synthesis of eye evolution by Nilsson (33, 34) departs in multiple ways from the traditional gradual-morphological model described in section 2a and therefore provides new opportunities to understand evolutionary origins during eye evolution. First, by making use of knowledge about how morphological structures interact with light, Nilsson’s synthesis explicitly considers function by calculating the amount of light required to perform different light-mediated behaviors. In earlier gradual-morphological models, functional considerations rarely went beyond the idealized notion that more intricate optical structures could be functionally ‘better’ and could evolve by natural selection. Second, Nilsson’s recent synthesis incorporates a more punctuated perspective than the gradual-morphological model by enumerating at least four structural innovations that underlie different organismal behaviors. After the origin of light sensing mechanisms, organisms with non-directional photoreception (Class 1) are able to sense changes in light intensity. After originating screening mechanisms such as light-absorbing pigments adjacent to photoreceptors, organisms gain directional photoreception (Class 2). After more finely dividing the visual field by adding a curved array of shielded photoreceptors, organisms may gain low-resolution spatial vision in dim light by adding specializations to the membranes of their photoreceptor cells that enhance photon capture (Class 3). Finally, after evolving a focusing apparatus, like a lens, organisms may evolve high-resolution vision (Class 4). In contrast, the earlier gradual-morphological models essentially considered eyes to lie along a gradual continuum from simple photoreceptive spot to camera eye (1). We hasten to add that although simple photoresponses may be achieved without nervous systems (35–37) complex eyes require rather elaborate nervous systems (e.g. 38). Although we could apply a similar approach to neural circuits, we restrict this review to the optical components of eyes.

Nilsson’s departures from the gradual-morphological model highlight a way forward for understanding origins in eye evolution. In particular, Nilsson’s synthesis defines critical innovations that each have implications for light sensitivity functions: phototransduction cascades, screening apparatuses, membrane elaborations, and focusing apparatuses. To these four, we add another innovation and hypothesize that a visual cycle - a specialized pathway for regenerating chromophores of visual pigments - may often be critical for vision that provides fine spatial and/or temporal resolution. Multiple of these innovations fit well into the component based approach we promote here because their genetic parts are characterized, allowing researchers to trace separately the origins of each of those parts. Next, we discuss what is known about each of these innovations, including definitions of each trait and its parts, estimates of each trait’s time of origin, and estimates of evolutionary histories and origins of the individual parts.

### a. Light detection: Origins of photosensitivity

The most basic light sensitivity function is a non-directional light sense. The simple detection of light is immediately useful for several organismal functions, including telling day from night, setting daily or seasonal rhythms, or determining depth in the water. Non-directional light sensors are often dispersed, found in the skin of many animals (39). The structural innovation required for light sensitivity is the molecular machinery for detecting light. Across all of life, there exist only a handful of different molecular mechanisms mediating biological responses to light, including Type II opsins (animal opsins), Type I opsins (e.g. halorhodopsin, bacteriorhodopsin, channelrhodopsins), lite-1, phototropin, neochrome, phytochrome, cryptochrome, light oxygen voltage (LOV) proteins and a photoactivated adenylyl cyclase (40–44). Although the component-based approach advocated here could be used to understand the origins of any of these molecular mechanisms of light sensitivity, here we focus on opsin-based phototransduction cascades because many components are well-studied and because they form the keystone functional modules of almost all animal eyes. We argue later that Type II and Type I opsins are not homologous, and that Type II opsins originated, within animals, in the common ancestor of eumetazoans. At the same time, other components of phototransduction are older, suggesting that opsin-based light sensitivity originated when the sensor in an older signal transduction pathway became sensitive to light.

#### Define the trait: Opsin-based phototransduction

Based on studies primarily in model species of flies and mammals, we have a good understanding of many parts of opsin-based phototransduction and their functional interactions. An important take-home message is that there is not a single, canonical phototransduction cascade, but rather at least four dramatically different ones (Table 1). Opsins are present in each cascade and are G-protein coupled receptors (GPCRs) that bind a light reactive chromophore (retinal). Collectively, an opsin/chromophore complex is called rhodopsin. When a photon strikes retinal, the chromophore changes from cis- to trans-, causing a concomitant change in the opsin protein. The opsin shape change exposes sections of the protein that activate signal transduction cascades leading to nervous impulses in the photoreceptor cells that house opsin. The non-opsin components of phototransduction are the components of GPCR pathways, including heterotrimeric G-proteins, intermediary enzymes, and ion channels. In addition to opsins from very divergent subfamilies, the non-opsin components vary and comprise different cascades.

**Table 1.** Four Different Phototransduction Cascades.

| Main model | G-alpha subunit | Intermediary Enzyme | Mechanism | Ion Channel |
| --- | --- | --- | --- | --- |
| Flies (156, 157) | G-α-q | Phospholipase C | Mechanical (158) | TRP |
| Vertebrates | G-α-t | Phosphodiesterase | cGMP hydrolysis | CNG |
| Scallops(159) | G-α-o | Guanylate cyclase | cGMP increase | K<sup>+</sup> channels |
| Box jellyfish (57) | G-α-s | Adenylyl cyclase | cAMP increase | CNG? (74) |

#### Estimate when opsin-based phototransduction originated

Estimating the origin of a trait involves comparing species’ phenotypes with their phylogenetic history. The trait of phototransduction can be conceived as a molecular phenotype, but it also is manifested as physiological phenotypes that can be scored as present or absent and compared to a phylogeny. Physiologically, phototransduction results in a change in a cell’s membrane potential that can be measured using electrical recordings from single cells. Even in the absence of single-cell recordings, electro-retinograms (ERG) and microspectrophotometry (MSP) may suggest opsin-based phototransduction. Comparing physiological phenotypes to animal phylogeny indicates that phototransduction is ancient because both bilaterian animals and some non-bilaterian animals have the trait (45, 46). This relative estimate for the timing of photoreceptor cell origins provides a framework for studying the components of these cells, especially the genes involved in opsin-based phototransduction. Did the components originate at the same time as the cells? If not, which components are new and which are old? What changes allowed the separate parts to function together? Separately addressing the evolutionary histories of the components can begin to illuminate these questions.

#### Determine when the components of opsin-based phototransduction originated

##### Opsins predate bilaterian animals

George Wald reported in the 1930s that the molecular basis of visual excitation in animals relies on a protein (now known as opsin) and a derivative of vitamin A (often retinal). And although scientists not long after had strong hints from protein biochemistry that homologous opsins are used in invertebrate and vertebrate vision, the first complete DNA and protein sequences of opsin genes were not determined until the 1980s (47–50). Based on these data, the homology of fly (a protostome) and cow (a deuterostome) opsin sequences clearly indicates that opsins originated at or before the first bilaterian animals (Fig.3).

**Figure 3.**
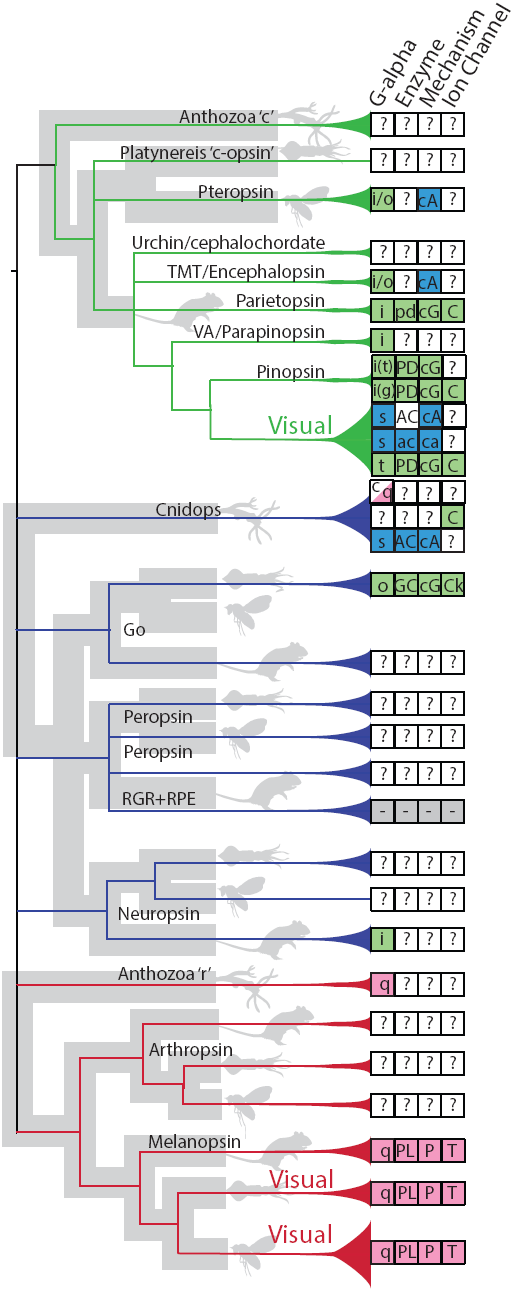
The evolution of opsins and phototransduction. Opsin gene-phylogeny is colored with green, blue and red branches. Grey species-phylogenies illustrate paralogous opsins in Cnidaria (hydra silhouette), Lophotrochozoa (squid silhouette), Ecdysozoa (fly silhouette), and Deuterostomia (mouse silhouette). The relationships shown are based mainly on Hering and Mayer (154), and the paralogous relationships implied - green as ‘c-opsins’, red as ‘r-opsins’, and blue as ‘type4’ opsins - are based on considerations from other analyses (59, 151). We exclude ctenophore opsins from this summary because of uncertainty in animal phylogeny and because ctenophore opsins fall close to known major groups (59, 154). At the right is a summary of functional experiments on phototransduction, focusing on cases where particular opsin genes are linked functionally to a G-proten alpha subunit, an intermediary enzyme, a cellular mechanism, and/or an ion channel. We colored components’ boxes green, red, or blue, based on their general association with Gi/o, Gq, or Gs alpha subunits. Abbreviations are as follows: o, t, g, i, s, q, c refer to different G-protein alpha subunits. t (=transducin), g (=gustducin), o, and i are all colored green because they are in one gene family. c refers to a cnidarian-specific G-protein (155). PL=Phospholipase C, PD=Phosphodiesterase (PDE) that increases cGMP, pd=PDE that lowers cGMP. AC=Adenylyl Cyclase, GC=Guanylate Cyclase. ca= cyclic AMP decrease; cA = cyclic AMP increase, cG = cyclic GMP, P=Phosphatidylinositiol 4,5-bisphosphate. C=Cyclic Nucleotide Gated (CNG) channel, Ck= a Potassium channel with properties similar to CNG. ‘-’ refers to isomerases that presumably do not initiate a GPCR cascade. Lophotrochozoa isomerases (retinochromes) are known, but they are highly diverged, making reliable phylogenetic placement difficult. We detail opsin groups and functional experiments from top to bottom of the figure, along with functional evidence, and references in Supplemental Table 1.

##### Opsins predate eumetazoan animals

Only more recently did clear evidence emerge that Type-II opsins exist outside bilaterian animals. Invertebrate zoologists knew that cnidarians and ctenophores possess photoreceptor cells similar to those of bilaterians, suggesting homology of cell type (51), and perhaps usage of opsins. In 2001, Musio et al. (52) found antibodies to squid opsin hybridized to ectodermal layers of *Hydra magnipapillata*, suggesting eyeless hydra use opsins. Finally, the determination of complete cnidarian genome sequences (53, 54) allowed Plachetzki et al. (55) to find cnidarian opsin homologs and characterize their expression in the eyeless *Hydra magnipapillata*. Shortly after, Suga et al reported expression of other cnidarian opsins (56). Koyanagi et al. (57) showed a cnidarian opsin to function in vitro as a visual pigment. Most recently, a ctenophore genome sequence confirmed that comb jellies possess opsins (58), and those genes fall in clades of already-known opsin groups (59). These studies clearly indicate that opsins originated at or prior to the eumetazoan ancestor (55). Furthermore, the absence of opsin in the genome sequence of the sponge *Amphimedon queenslandica*, the choanoflagellate *Monosiga brevicollis*, and multiple fungi, including *Saccharomyces cerevisiae*, suggested Type II opsins might have originated within animals (55).

##### Opsins do not predate animals

Although there exist some Type-II-opsin-like sequences outside of animals, there is no evidence that those are functional opsins. One intriguing candidate was identified with genomic sequencing: a Type-II-opsin-like sequence in a fungus Based on sequence similarity, Heintzen (60) reported an opsin-like gene in the chytrid fungus *Allomyces macrogynus*, a particularly intriguing result in light of an earlier study on a related species. In *Allomyces reticulatus*, Saranak and Foster (61) interrupted retinal synthesis to demonstrate that the phototactic behavior of zoospores depends on retinal. Because opsins (Type I and Type II) are the only known retinylidene proteins (40, 41), the use of retinal in phototaxis indicated *A. reticulatus* very likely uses opsin to detect light. But recent experiments on phototactic zoospores of a third closely related species (*Blastocladiella emersonii*) indicate that light sensitivity is not due to a Type-II-like opsin, but rather to a fusion gene that includes a Type I opsin (62). Because most chytrid Type-II-like opsins lack a lysine at a position homologous to the Schiff base of known opsins, and because phototactic behavior can be ascribed to a Type-I opsin, there remains no convincing evidence for Type-II opsins outside of animals. Therefore, we maintain that Type-II opsins originated near the common ancestor of eumetazoan animals (55).

Some researchers suggest that Type-I and Type-II opsins are homologous, and since Type-I are present in bacteria, this would point to a very ancient origin of opsins. Mackin et al. (63) recently resurrected this homology hypothesis, claiming structural similarities and a lack of functional constraint indicate homology. They further speculate that Type-I opsins evolved “from Type-II opsins” through an ancient horizontal transfer event that led to erasure of all historical signal of homology, except for primary structural features (63). While that hypothesis is challenging to test scientifically, a number of facts point instead to non-homology and therefore convergent origins of Type-I and Type-II opsins. First, there is no amino acid similarity beyond that expected of random sequences (41). Second, crystal structures are known from Type-I and Type-II opsins and structural comparisons show significant differences in the size and organization not only of hydrophilic regions, but also in the arrangement of seven transmembrane domains (reviewed in 41). Third, only Type-II opsins interact with G-proteins. Fourth, Type-I opsins show a signal of origin by domain duplication, whereas Type-II opsins show no such signal (64). Fifth, Type-I and Type-II opsins often utilize retinal chromophores with different chemistries (reviewed in 41). This evidence indicates the similarities between Type-I and Type-II opsins evolved convergently, consistent with our assertion that Type-II opsins originated within animals.

#### Other phototransduction components predate animals

Phototransduction cascades are typical GPCR cascades, and while opsins originated within animals from non-light sensitive receptors, other components of phototransduction have more ancient origins. Therefore, the origin of the first animal phototransduction cascade probably evolved when a GPCR became light sensitive, perhaps by evolving to bind a retinal chromophore as all extant opsins do today. The additional diversity of phototransduction cascades evolved later, again from existing components. Here, downstream parts - including arrestins, G-proteins, intermediary enzymes, and ion channels - probably were used in other GPCR cascades before their functional association with opsin. If so, their ancient histories indicate that duplicated opsins diversified by co-opting existing downstream components of other GPCR pathways to form multiple phototransduction cascades, probably caused by mutations in opsin that activated different G-proteins (4, 55, 65).

Phototransduction components that predate phototransduction itself include (at least): arrestins, G-protein-alpha subunits, and CNG and TRP ion channels. Arrestins function in signal transduction by interacting with GPCRs like opsin to regulate receptor activity (reviewed in 66). Functionally characterized arrestins are present in deuterostomes (67) and protostomes (68, 69), and sequences are known from sponges and choanoflagellates (70). Core arrestin domains are present in fungi in pH sensors (71). Opsins activate heterotrimeric G-proteins, which also predate animals, as they are known from fungi and plants (72). Even the different G-protein subunit genes that help define different phototransduction cascades (G-□-s, G-□-t, G-□-o, and G-□-q) originated by duplication prior to metazoans (73). Cyclic Nucleotide Gated (CNG) ion channels are involved in a variety of signaling cascades and may be the ancestral ion channel in phototransduction (74). If so, signaling cascades involving TRP probably were co-opted somewhat later (74), although still early in animal evolution. TRP ion channels form an ancient, and structurally and functionally diverse superfamily, with those involved in phototransduction being from the TRPC subfamily, which itself predates animals (reviewed in 75).

## b. Screening pigments: Multiple origins of directional photoreception

By combining screening pigments and photoreceptors together to evolve eyespot structures, organisms gain the function of directional photoreception, useful for tasks such as phototaxis and the detection of motion (34). Eyespots can be quite simple morphologically, yet even single photoreceptors may provide directional photoreception to mobile organisms if they are screened by pigment appropriately (36, 76). In more complex visual organs, screening pigments help preserve the contrast of images through the absorption of off-axis photons and they may contribute to dynamic range control. Therefore, screening pigments not only aid visual function in animals that lack image-forming eyes, but also retain their utility if additional optical refinements arise. The evolution of screening pigments - especially their synthesis - can be subjected to a component-based study because many of the genes responsible for generating pigments are deeply conserved and many have been well characterized in model species (77– 81). Even though the proteins comprising synthesis pathways of pigments are deeply conserved, the mechanisms for patterning pigments evolve more rapidly through changes in expression of synthesis pathway genes (e.g. 82). Studying the deep evolutionary history of changes in gene expression is more challenging than comparing protein similarity and phylogenetic relationships. Therefore, understanding the deep history of pigment synthesis is more approachable than the deep evolution of pigment patterning, even though both are required for a full understanding of eye evolution.

### Define the trait: Screening pigments

Here, we focus on the evolutionary histories of the three types of pigment most commonly used by metazoans to screen the photoreceptors of their eyes and eyespots: melanins, pterins, and ommochromes (83). Melanins, synthesized from the amino acid tyrosine, include the yellow to red-brown pheomelanins and the brown to black eumelanins (84)(85). Pterins (also called pteridines) are synthesized from purines, such as guanosine triphosphate (GTP), and range in color from yellow to orange to red (86). Ommochromes - which tend to be yellow, red, brown, or black - are derived from the amino acid tryptophan (87)(88). Certain organisms screen their photoreceptors in other ways, but these tend to be either too generalized in form or too rare to be studied through a comparative, component-based molecular approach (89–91). While pigments may be guessed from color, to distinguish conclusively between different types of screening pigments, researchers test solubility. Melanin is insoluble, but pheomelanins, pterins, and ommochromes tend to be soluble in sodium hydroxide, ammonium hydroxide, and acidified methanol, respectively. Once pigments are extracted, researchers characterize absorption spectra and use techniques like HPLC and mass spectroscopy to verify identity. These techniques are reliable because readily available standards can be compared to naturally-occurring pigments (or their degradation products) e.g (92, 93).

### Estimate when screening pigments originated in separate lineages

While it is clear that melanin, pterin, and ommochrome pigment synthesis has deep evolutionary origins, we know little about when particular pigments became associated with particular visual organs (but see 83). To estimate when screening pigments were co-opted within lineages as components of particular visual systems, we will need to know more about the specific taxonomic distributions of pigments and to build phylogenies with sampling strategies appropriate for learning when, where, and how many times certain types of screening pigments became functionally associated with photoreception. Yet it is already clear that melanins, pterins, and ommochromes are widespread taxonomically and that all three types of pigment tend to be used by organisms for a variety of tasks. Melanins (e.g. 82), pterins (94, 95), and ommochromes (96, 97) all benefit organisms by creating color patterns useful for camouflage and signaling (98). Melanins and pterins also may help prevent oxidative damage to organisms by absorbing UV radiation and scavenging free radicals (reviewed by (99)). From these putatively more ancestral roles (because they are so widespread phylogenetically), it is likely that melanins, pterins, and ommochromes were each co-opted separately by several different lineages of animals to act as screening pigments for their eyespots or eyes. For example, melanin-based screening pigments have been identified in certain cnidarians, turbellarian flatworms, nematodes, polychaetes, molluscs, non-vertebrate chordates, and vertebrates (reviewed by 83). Pterins are associated with the eyes of certain arthropods (100) and annelids (101) and ommochromes are expressed by the compound eyes of certain arthropods (102), as well as the camera eyes of cephalopod mollusks (103). Placing these data on pigment types in photoreceptors in an explicitly phylogenetic comparative context, and especially identifying particular transitions from photoreceptor to eye spot, will yield more specific hypotheses about when screening pigments became associated with other components of photoreceptors.

### Determine when the genetic components of pigment synthesis originated

#### The synthesis pathways for melanins, pterins, and ommochromes are deeply conserved

From a phylogenetic perspective, the genetic potential to produce screening pigments tends to precede the origins of eyespots or eyes in lineages. That is, many metazoans lack pigmented eyespots or eyes, but nearly all taxa either produce melanins, pterins, and/or ommochromes or have the genetic potential to produce these pigments. For example, the vast majority of melanin-expressing organisms synthesize melanins by using the enzyme tyrosinase (TYR), a monophenol oxygenase, to convert tyrosine to DOPA (3,4-dihydroxyphenylalanine) and DOPA to DOPAquinone (84, 85). Given evidence that TYR helps produce melanin in fungi (104) and that this enzyme can produce melanin from tyrosine without the involvement of other enzymes, it appears that TYR has a deep evolutionary history as a producer of melanin (78, 105). Arthropods appear to be the one exception to this pattern, as they produce melanin through a synthesis pathway in which the enzymes phenoloxidase (PO) and dopachrome conversion enzyme (DCE) assume the functional roles of TYR and TRP2, respectively (77)(106, 107). Phenoloxidases, close relatives of the hemocyanins of arthropods, originated early in the evolutionary history of arthropods (108) and it appears that DCE also represents an arthropod-specific invention (109).

As in the case of melanins, it is likely that the common ancestor of Metazoa had the capacity to produce both pterins and ommochromes. Pterin synthesis involves the conversion of the purine GTP to H_4_biopterin through the enzymatic actions of GTP cyclohydrolase 1 (GCH1), 6-pyruvoyl H4pterin synthase (PTS), and sepiapterin reductase (SPR) (100, 105). All three of these enzymes originated prior to the appearance of metazoans: GCH1, PTS, and SPR contribute to the synthesis of H_4_biopterin in the fungus *Mortierella alpine* (110) and phylogenetic evidence suggests that homologs of these genes may be present in bacteria (78). The components of the ommochrome synthesis pathway include tryptophan 2,3-dioxygenase (TDO2), kynurenine formamidase (KF), and kynurenine 3-monooxygenase (KMO) (87, 88, 100), all of which are deeply conserved among metazoans (78) and beyond (*e.g.* KF is found in yeast (111)).

## c. Membrane elaborations: Enhanced sensitivity for vision in low light

Another critical evolutionary innovation with important functional consequences is the elaboration of the membranes of photoreceptor cells. When the membranes of photoreceptor cells are elaborated - for example by invaginations that lead to stacking - the probability that the cell will capture a photon increases because more rhodopsin molecules can be packed in the increased surface area of the cell. Therefore, membrane elaboration increases the efficiency of photoreceptors, allowing for visual organs that are able to operate at lower levels of light. The increased efficiency of photon capture may be required for eyes that allow directional light sensitivity and high performance vision because those eyes split a field of view into fine subdivisions, and there are a finite number of photons passing through any given field. When and how did membrane elaborations of photoreceptors evolve?

### Define the trait: Membrane elaboration

The lure of a simple two-category classification for photoreceptor cells based on membrane elaborations has both inspired biologists and led to controversies. Eakin pioneered electron microscopy techniques, which remain the best way to characterize photoreceptor morphology and he proposed a two-category classification (112). In Eakin’s classification, photoreceptor cells either evolved increased surface area by elaborating a cilium or adding numerous cilia (ciliary) or by adding numerous invaginations (rhabdomeric) to the cell membrane. Salvini-Plawen and Mayr (1) suggested two additional categories, but few biologists follow their lead today. The advent of molecular data reinvigorated Eakin’s classification because ciliary receptors tend to use hyperpolarizing phototransduction and a particular clade of opsins, while rhabdomeric receptors tend to use depolarizing, TRP-based phototransduction and another clade of opsins (113). Unfortunately, there exist exceptions not only to clear morphological classification of cell types (114), but also to a simple relationship between phototransduction and cell type (115).

### Determine when membrane elaborations originated

Membrane elaborations in photoreceptor cells include microvilli and cilia, which also have evolutionary histories older than photoreceptors. Microvilli are comprised of cross-linked actin filaments and are present not only in animals, but also in choanoflagellates (116). Furthermore, microvilli are structurally related to filopodia, which have an even more widespread phylogenetic distribution (116). Given their varied function and broad phylogenetic distribution, microvilli very likely evolved before phototransduction in a different functional context. Cilia, and structurally related organelles like flagella, have a microtubule-based cytoskeleton and are present in the cells of most eukaryotes (117, 118). Interestingly, the cAMP and cGMP signalling mechanisms used in photoreceptor cells are also present and associated with cilia in most eukaryotes, suggesting an ancestral role of cyclic nucleotide signaling in sensory cilia (119, 120). Even though cilia were lost secondarily in several lineages, like plants and most fungi, they play a dominant role in animal sensory structures. Similar to microvilli, cilia have varied function and a broad phylogenetic distribution indicating they too evolved before phototransduction.

### Determine when the components of membrane elaboration phototransduction

A component-based approach very similar to what we outline here for eyes has been published for both microvilli (116) and cilia (117, 118). The components of these organelles extend much earlier than animals, and much earlier than their use in photoreceptors. Since microvilli, cilia, and their components originated long before animals, the key information for the evolution of photoreceptors and eyes is when the organelles became functionally associated with other photoreceptive components. Here, we still have much to learn.

## d. Image formation mechanisms: Multiple transitions to high-resolution vision

Image formation (focusing) mechanisms - such as lenses or mirrors - improve the resolution of eyes by restricting the angular regions of space from which individual photoreceptors gather photons (121). Thus, by adding a focusing mechanism to a collection of pigmented photoreceptors, an animal will be able to gather finer-grained spatial information from its environment. As an additional benefit, focusing mechanisms help alleviate trade-offs between resolution and sensitivity (121): the sensitivity of a pigment-cup eye is proportional to the cross-sectional areas of its photoreceptors (a matter of microns), but the sensitivity of an eye with a lens is proportional to the cross-sectional area of the lens (a matter of millimeters or centimeters) (34). Additionally, the origins of focusing mechanisms may be relatively simple from a molecular perspective. The evolution of an imaging lens, for example, may begin with the expression of any water-soluble protein resistant to light-induced damage or self-aggregation; as long as such a proto-lens has a refractive index higher than the surrounding medium, it will help improve both the resolution and sensitivity of an eye (121). With these predictions in mind, it is not surprising that focusing mechanisms evolved multiple times in animals through the co-option of a wide range of molecular components.

### Define the trait: Focusing structures

#### Animals may use lenses, corneas, and/or mirrors as focusing mechanisms

The eyes of most animals form images using light-refracting structures such lenses and/or corneas. Biological lenses are relatively thick structures that lie below an animal’s epithelium or cuticle. Compared to lenses, corneas are relatively thin structures that are formed from modified regions of the outermost layer of an animal’s body. Aquatic organisms generally use lenses as focusing structures because lenses tend to provide more refractive power than corneas. Terrestrial organisms - such as vertebrates and spiders - tend to use their corneas for image formation, a workable option because corneas provide more focusing power in air than in water (122). The vast majority of biological lenses and corneas are composed of proteins (19), but several distantly related lineages of animals have eyes with focusing structures made of minerals such as calcium carbonate (CaCO_3_) (e.g. 123). As an alternative to lenses or corneas, certain animals use mirrors to form images by reflection instead of refraction (124).

### Estimate when focusing mechanisms originated in separate lineages

#### Focusing mechanisms evolved multiple times in animals

Although there is some disagreement over the number of times that image-forming eyes (*i.e.* eyes with focusing mechanisms) evolved in metazoans, authors generally list over a dozen separate occurrences (1, 25, 125). Many of these eyes are superficially similar, but phylogenetic and structural analyses indicate that their image-forming capabilities evolved separately. For example, camera-type eyes with proteinaceous lenses are found in cubozoan cnidarians, cephalopod mollusks, and vertebrates. The proteins that compose the lenses of these eyes are non-homologous, supporting separate instances of the evolution of spatial vision (19). A phylogenetic perspective offers further evidence that camera eyes with lenses have evolved separately in different lineages. Cubozoans, for example, are in a derived position within Cnidaria (126) and most other cnidarians lack lensed eyes. Similar arguments may be applied to other eyes as well, allowing researchers to determine when and how many times certain focusing mechanisms evolved. Given what appear to be limited phylogenetic distributions, it is possible that mineral-based lenses and image-forming mirrors represent relatively recent appearances of spatial vision among animals.

### Determine when the genetic components of focusing structures originated

#### Lenses evolve by co-opting stress-response proteins

Although we can take a similar component-based approach to studying the origins of corneas, mineral-lenses, and image-forming mirrors, we here focus on biological lenses, which tend to derive refractive power from expressing high concentrations of water-soluble proteins termed crystallins (19). Crystallins have evolved separately in different lineages of animals and they often originate from proteins expressed broadly outside of eyes with ancestral roles in stress responses (19). For example, the crystallins expressed in the lenses of vertebrates are small heat shock proteins (127) that are up-regulated in response to stress in a broad range of tissues outside of eyes (128, 129). Similarly, the diverse S-crystallins that provide focusing power in the separately evolved camera eyes of cephalopods are derived from glutathione-S-transferase (130), a stress-response protein (and, perhaps not coincidentally, also a member of the pterin pigment-synthesis pathway (131)). As a final example, the omega-crystallins of cephalopods (132, 133) and scallops (134) are aldehyde dehydrogenases, representing two separate cases where a third class of stress response enzyme was co-opted as a lens protein. Thus, it appears that the molecular components required for producing lenses in animals tend to have origins that greatly precede those of the lenses themselves. As a counter-example, the lenses of the adult eyes of the cubozoan cnidarian *Tripedalia cystophora* express cnidarian-specific proteins known as J-crystallins (130). Here is a case, intriguing in its rarity, where the timing of the origin of new genes matches closely with the appearance of a new focusing structure.

## e. Visual cycles: Origins of high-performance vision?

Opsins bind retinal, a derivative of Vitamin A that undergoes a *cis-*to-*trans* isomerization when it absorbs a photon of light (115, 135). For retinal to be re-used by an opsin for light detection, it must be converted back to its *cis* form (136). Some animals use specialized enzymatic pathways - known as visual or retinoid cycles - to convert *trans*-retinal to *cis*-retinal. Following previous authors, we hypothesize that visual cycles evolved multiple times in Metazoa (137–139). We hypothesize that visual cycles co-evolve with image-forming eyes that provide fine spatial and temporal resolution. High-performance eyes require large amounts of *cis*-retinal to be supplied at rapid rates because fine spatial resolution requires large numbers of photoreceptors and fine temporal resolution requires photoreceptors that are highly sensitive to light (34). Alternately, visual cycles may evolve as mechanisms for fine-tuning the sensitivities of photoreceptors that operate in variable light environments. For example, a visual cycle could tune the sensitivities of photoreceptors by supplying large amounts of *cis*-retinal when light levels are low and relatively small amounts of *cis-*retinal when light levels are high.

### Define the trait: Visual cycles

#### Do visual cycles and high-performance eyes co-evolve?

Visual cycles are multi-step, enzymatic pathways that convert *trans*-retinal to *cis-*retinal for the use of opsins expressed by photoreceptor cells (140, 141). Here, we distinguish visual cycles from photo-reconversion (also termed photo-regeneration), a phenomenon in certain bistable visual pigments in which *trans*-retinal remains bound to an opsin and is converted back to *cis-*retinal when it absorbs a photon of an appropriate wavelength. As in our inquiries into the molecular components of phototransduction and pigment synthesis, characterizing the components of visual cycles and studying their distributions and modes of origin will help us learn how and why visual cycles have evolved in different lineages. For example, we can test whether visual cycles and high-performance eyes tend to co-evolve, perhaps due to the latter’s need for large amounts of *cis-*retinal. Following past reports (e.g. 137), we can also explore whether the visual cycles of animals employ lineage-specific genes with evolutionary histories that are shallow relative to those of other genetic components of photosensitivity and eyes.

### Estimate when and how many times visual cycles evolved

#### Visual cycles differ between lineages

Lineages of animals differ in the molecular mechanisms that they use to convert *trans*-retinal to *cis*-retinal. Specifically, recent discoveries reveal that the visual cycles of vertebrates (140), cephalopod mollusks (142), and fruit flies (139) differ in fundamental ways. The Gt-coupled opsins of vertebrates, for example, release retinal once it absorbs a photon of light. The reconversion of *trans*-retinal to its *cis-* form subsequently involves a multi-step, intercellular process that includes a specialized form of opsin termed RGR (143). Cephalopod mollusks use an opsin closely related to RGR - termed retinochrome - for their intracellular visual cycle; however, the other components of this pathway differ from those of vertebrates (142). Until recently, it was thought that a visual cycle was absent in arthropods because the Gq-couple opsins expressed in their eyes are bistable and thus capable of photo-reconversion. We now know that *Drosophila* has a distinct, multi-step visual cycle and that the components of this cycle are required for normal visual performance (141). A gap in our current knowledge is whether visual cycles are present in the low-resolution eyes or eyespots found in many animals. From a Metazoa-wide perspective, the eyes of vertebrates, cephalopod mollusks, and insects tend to provide relatively fine spatial and/or temporal resolution. To search for correlations between the origins of high-performance eyes and visual cycles, we will need to learn whether visual cycles are present or absent in simple eyes that require comparatively small amounts of *cis*-retinal to operate.

### Estimate when the molecular components of visual cycles originated

#### Visual cycles employ lineage-specific genes

The camera eyes of vertebrates and cephalopods use specialized and evolutionarily related forms of opsin - termed RGR (143) and retinochrome (142) - to recycle retinal by converting it from *trans-* to *cis-* conformation. Aside from their use of related opsins as photoisomerases, the visual cycles of vertebrates and cephalopods employ dissimilar genetic components. The evolutionary origins of these components tend to be recent relative to the origins of the molecular components of opsin-based phototransduction and pigment synthesis. For example, the visual cycle of vertebrates likely originated in the common ancestor of jawed and jawless fish. Two genes critical to the vertebrate visual cycle – lecithin:retinol acyltransferase (LRAT) and *all-trans* retinyl ester isomerohydrolase (RPE65) – are present and capable of aiding in the visual cycle of lampreys (*Petromyzon marinus*), but are either absent or incapable of appropriate enzymatic activity in non-vertebrate chordates (144). Given evidence that the common ancestor of lampreys and jawed vertebrates possessed a complex camera eye (145), it is possible that high-performance vision and a visual cycle co-evolved in vertebrates. Like vertebrates, cephalopods employ lineage-specific components in their visual cycle. Specifically, cephalopods use the retinal-binding protein RALBP to transport retinal between rhodopsin and retinochrome (146, 147). The phylogenetic distribution of RALBP outside of cephalopods remains largely unexplored, but there is evidence that a visual cycle based on RALBP and retinochrome may be present in the eyes of other mollusks as well (148).

Despite employing Gq-coupled opsins that are bistable, the eyes of arthropods also show evidence of separately evolved visual cycles. Recently, Wang et al. (141) compiled evidence that *Drosophila melanogaster* has an enzymatic visual cycle that functions separately from the *de novo* synthesis of retinal. Enzymes involved in *Drosophila*’s visual cycle that are necessary for normal visual function include a pigment-cell-enriched dehydrogenase (PDH) (139) and a retinol dehydrogenase (RDHB) (141). Further, the eyes of certain butterflies (*e.g. Papilio xuthus*) express retinol-binding proteins that are not observed outside the superfamily (149, 150). It is likely that additional components of the visual cycles of fruit flies, butterflies, and other arthropods have yet to be discovered.

## 6. Synthesis

The history of eyes and the genes involved is a history of tinkering and bricolage. Although eye-specific genes exist in a few particular animal clades, practically every gene family expressed in eyes contains genes that function outside of eyes and that have ancient histories that predate animals. Therefore, the genes of today’s eyes had other uses before they formed new functional associations. Genes of eyes combined forces at different times, in different clades, and in different combinations, to detect, screen, and/or focus light. We notice that many of these gene families that function in eyes may have ancestral roles in stress response, pointing to a possible mechanism to influence co-option. Examples include opsin genes, which are related to melatonin receptors (74, 151). Melatonin breaks down in light and may have originated as an anti-oxidant (152). Additionally, pigments are often used to protect cells from light. Genes produce melanin and pterin pigments, which absorb potentially harmful photons and reduce tissue damage by acting as anti-oxidants (99). Different genes for lens proteins are also stress enzymes. For example, α β -crystallin/HSP has a role in eyes to protect mitochondrial function against oxidative stress (153). Because light causes photo-oxidative stress and DNA damage, protective responses are mounted in cells exposed to much light. We speculate that during evolutionary history, multiple genes that now interact for visual function, originated in stress responses to light.

Our study of eye evolution inspires a research program to break complex traits into component parts to understand the timing and mechanism of their origins. This approach may be applied to most any biological trait, and when applied broadly, will allow insights into fundamental evolutionary questions. Do interacting parts originate gradually or abruptly? What types of mutation more commonly lead to trait origins? What changes first during evolution, structure or function? What are the environmental contexts of trait origins? The answers to these questions will enrich our understanding of the evolutionary process, and continue to provide clear, scientific explanations for how biological complexity has originated.

## Acknowledgements

We thank D-E Nilsson, M Porter, D Plachetzki, T Cronin, and Oakley’s lab for comments on earlier drafts. This work was supported by grants from the National Science Foundation.

---

**Table S1.**
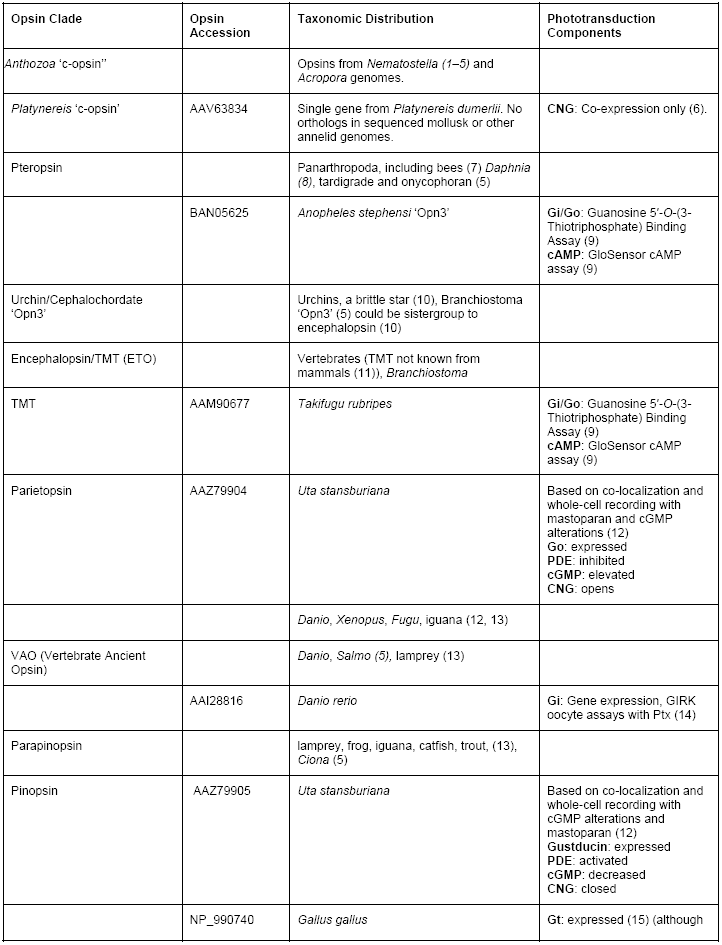
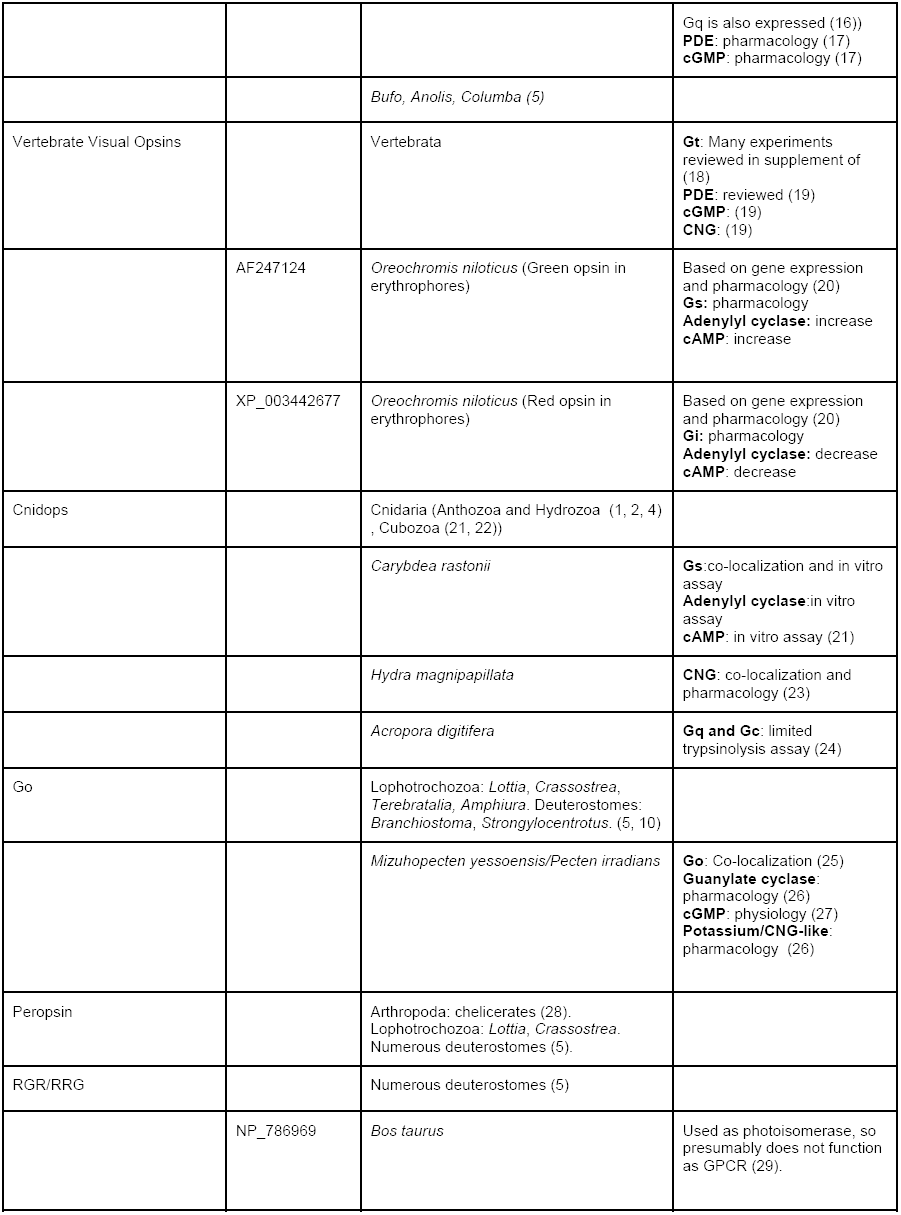
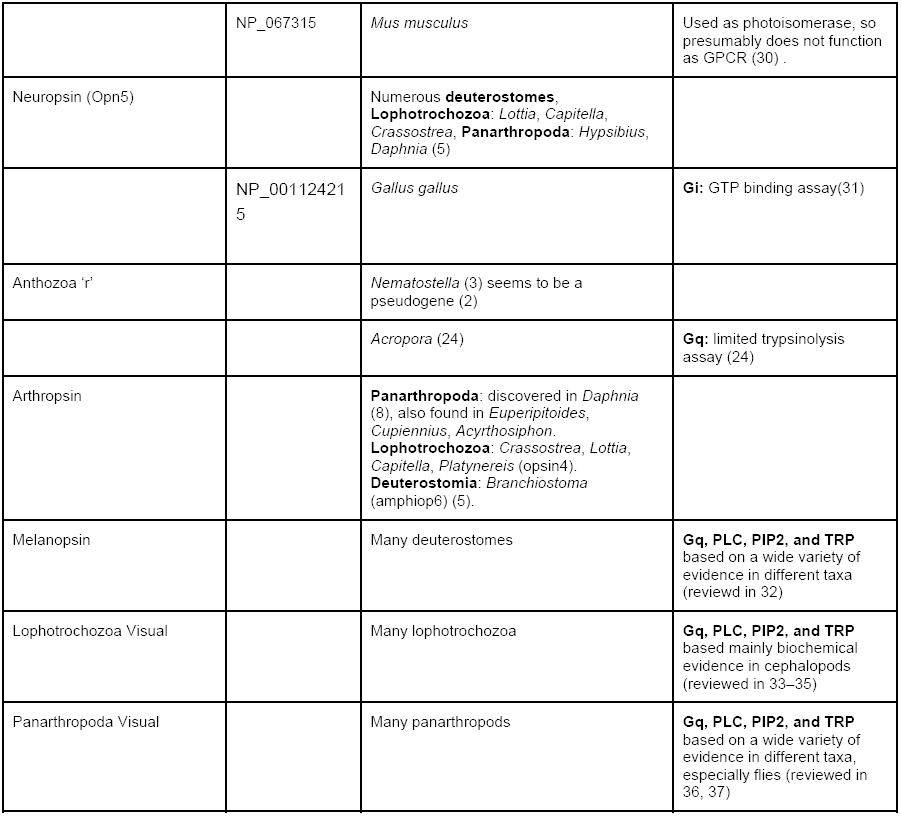
Summary of opsins, opsin clades, and functional information about phototransduction components.

